# The readability of scientific texts is decreasing over time

**DOI:** 10.1101/119370

**Authors:** Pontus Plavén-Sigray, Granville James Matheson, Björn Christian Schiffler, William Hedley Thompson

## Abstract

Clarity and accuracy of reporting are fundamental to the scientific process. The understandability of written language can be estimated using readability formulae. Here, in a corpus consisting of 707 452 scientific abstracts published between 1881 and 2015 from 122 influential biomedical journals, we show that the readability of science is steadily decreasing. Further, we demonstrate that this trend is indicative of a growing usage of general scientific jargon. These results are concerning for scientists and for the wider public, as they impact both the reproducibility and accessibility of research findings.

## INTRODUCTION

Reporting science clearly and accurately is a fundamental part of the scientific process, facilitating both the dissemination of knowledge and reproducibility of results. The clarity of written language can be quantified using readability formulae, which are well-established estimates of reader understandability (1, 2, 3, 4, 5, 6). Texts written at different times can vary in readability: trends towards simpler language have been observed in US presidential speeches (7), novels (8, 9) and news articles (10). There are studies that have investigated linguistic trends within the scientific literature. One study showed an increase in positive sentiment (11), finding that positive words such as “novel” have increased dramatically in scientific texts since the 1970s. A tentative increase in complexity has been reported in scientific texts in a limited dataset (12), but the extent of this phenomenon and any underlying reasons for such a trend remain unknown.

To investigate trends in scientific readability over time, we downloaded 707 452 article abstracts from PubMed, from 122 high-impact journals selected from twelve biomedical fields of research (Fig. 1A-C). This journal list included, among others, Nature, Science, NEJM, Lancet, PNAS and JAMA (see Supplementary Methods and Supplementary Materials S1) and ranged from 1881 to 2015. We quantified the reading level of each abstract using two established measures of readability: the Flesch Reading Ease (FRE (1, 2)) and the New Dale-Chall Readability Formula (NDC, (3)). The FRE is proportional to the number of syllables per word and the number of words in each sentence. The NDC is proportional to the number of words in each sentence and the percentage of ‘difficult words’. Difficult words are defined as those words which do not belong to a predefined list of common words (see Methods). Lower readability is indicated by a low FRE score or a high NDC score (Fig. 1A).

**Figure 1:**
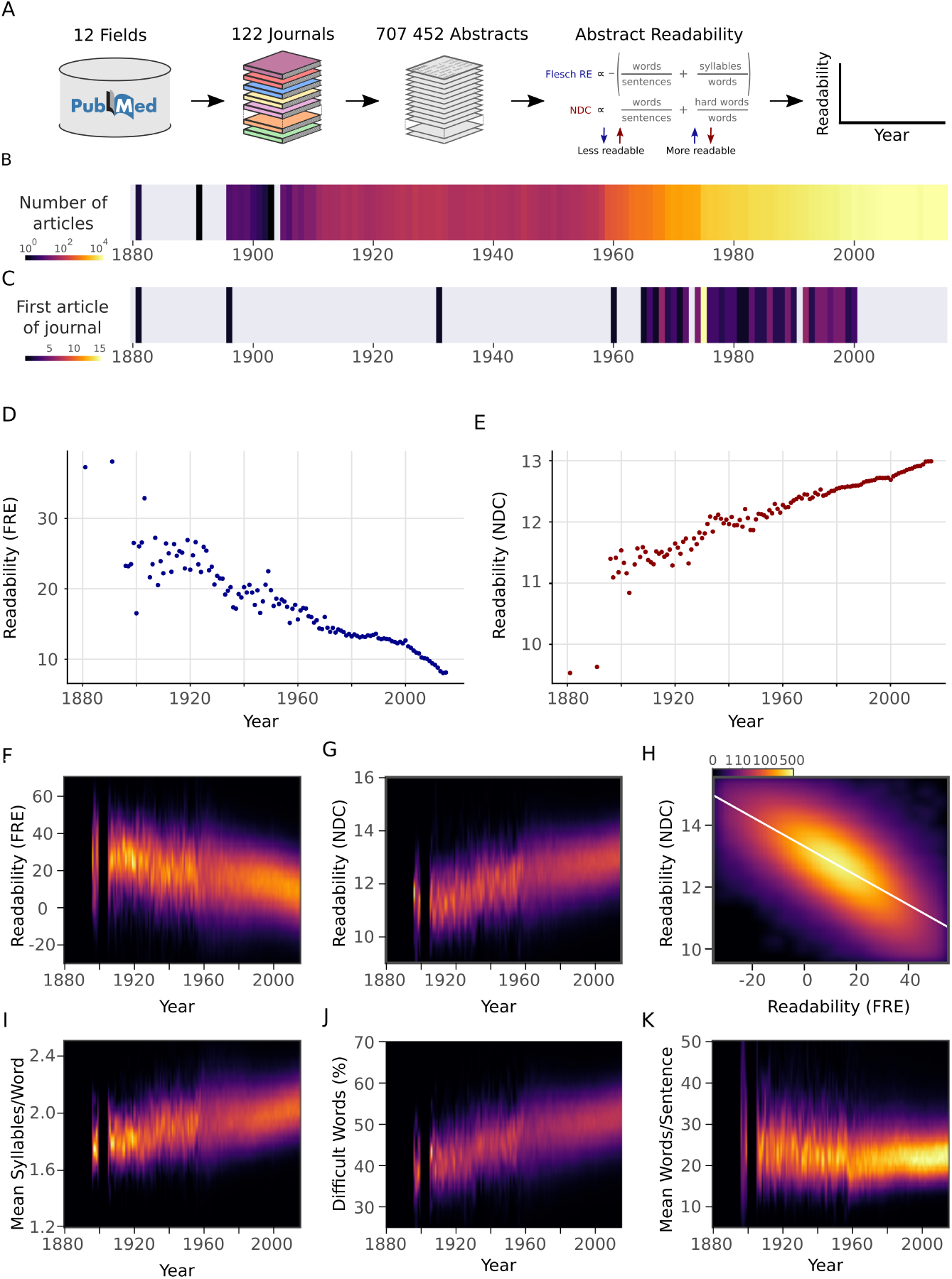
Readability of scientific abstracts decreases over time. **a**, Schematic depicting the major steps in the abstract extraction and analysis pipeline. **b**, Number of articles in the corpus published in each year. Colour scale is logarithmic. **c**, Starting year of each journal within the corpus. This corresponds to the first article in PubMed with an abstract. **d**, Mean Flesch Reading Ease (FRE) readability for each year. Lower scores indicate less readability. **e**, Mean New Dale-Chall (NDC) readability for each year. Higher scores indicate less readability. **f**,**g**, Kernel density estimates displaying the readability (f: FRE, g: NDC) distribution of all abstracts for each year. **h**, Relationship between FRE and NDC scores across all abstracts, depicted by a two-dimensional kernel density estimate. Axis limits are set to include at least 99% of the data. Colour scales are exponential. **i**-**k**, Kernel density estimates displaying the components of the readability measures (i: syllable to word ratio; j: percentage of difficult words; k: word to sentence ratio) distribution of all abstracts for each year. For kernel density plots over time (f, g, i, j, k), years with fewer than ten abstracts are excluded to obtain accurate density estimates. Lighter colours depict more abstracts.

## RESULTS

The primary research question was to examine the relationship between article abstract readability with year of publication. We observed a strong decreasing trend of the average yearly FRE (r = -0.93, *p* < 10^−15^) and a strong increasing trend of average yearly NDC (r = 0.93, *p* < 10^−15^) (Fig. 1D-H). Next, we examined the relationship between the components of the readability metrics over years. The average number of syllables in each word (FRE component) and the percentage of difficult words (NDC component) showed pronounced increases over years (Fig. 1I,J). Sentence length (FRE and NDC component) showed a steady increase with year after 1960 (Fig. 1K), the period in which the majority of abstracts were published (Fig. 1B). FRE and NDC were correlated with one another (r = -0.72, *p* < 10^−15^) (Fig. 1H).

The readability of individual abstracts was formally evaluated in relation to year of publication using a linear mixed effects model with journal as a random effect for both measures. The fixed effect of year was significant (FRE: *b* = -0.19, *p* < 10^−15^; NDC: *b* = 0.016, *p* < 10^−15^; Supplementary Materials S2). The average yearly trends combined with this statistical model reveal that the complexity of scientific writing is increasing with time.

To verify that the readability of abstracts was representative of the readability of the entire articles, we downloaded full text articles from six additional independent journals from which all articles were available from the PubMed Central Open Access Subset (Fig. 2A). Although, as has previously been reported (13), abstracts are less readable than the full articles, there was a strong positive relationship between readability of the abstracts and the full texts (FRE: *r* = 0.58, *p* < 10^−15^; NDC: *r* = 0.63, *p* < 10^−15^ Fig. 2B, Supplementary Materials S3). This implies that the increasing complexity of scientific writing generalises to the full texts.

**Figure 2:**
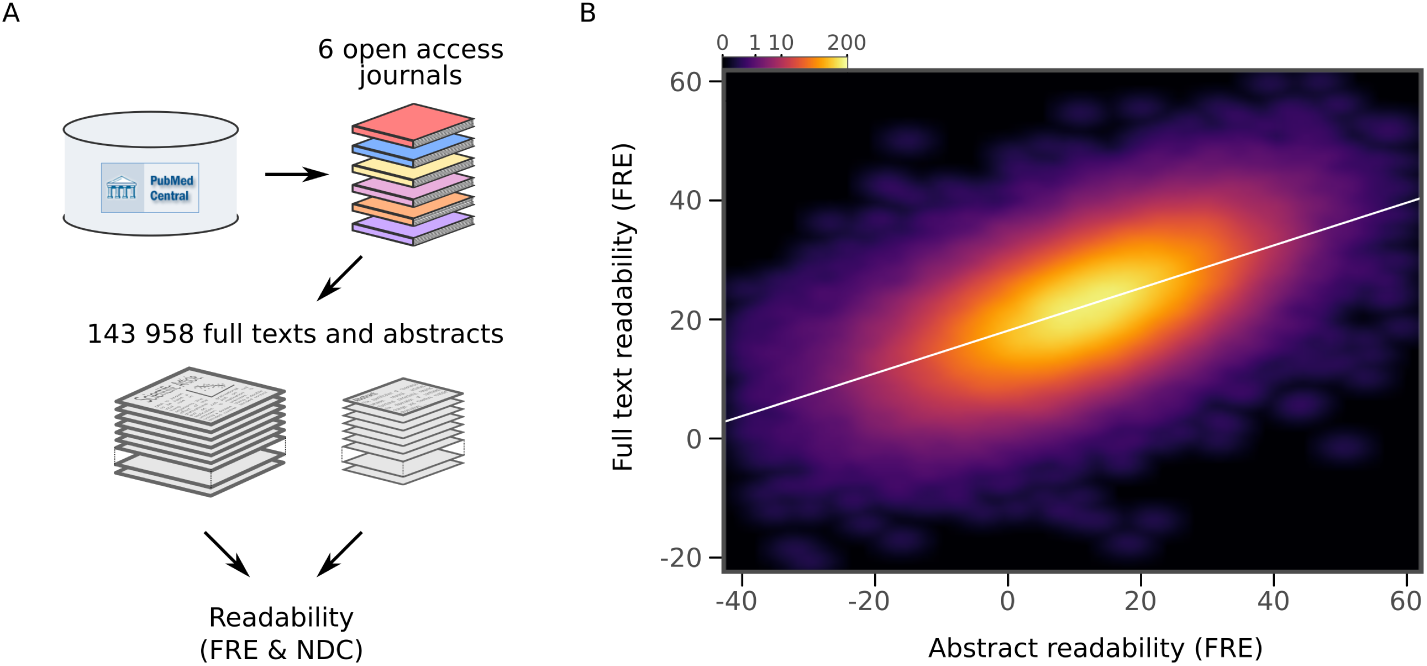
Readability of scientific abstracts correlates with readability of full texts. **a**, Schematic depicting the major steps in the full text extraction and analysis pipeline. **b**, Relationship between Flesch Reading Ease (FRE) scores of abstracts and full texts across the full text corpus, depicted by a twodimensional kernel density estimate. Axis limits are set to include at least 99% of the data. For New Dale-Chall (NDC), see Supplementary Materials S3.

There could be a number of explanations for the observed trend in scientific readability. We formulated two plausible and testable hypotheses: (1) There is an increase in the number of co-authors over time (Fig. 3A) (see also 14, 15). If the number of co-authors correlates with readability, this underlies the observed effect (i.e. a case of ‘too many cooks spoil the broth’). (2) An increase in a general scientific jargon is leading to an in-group vocabulary which is less readable (i.e. a ‘science-ese’).

**Figure 3:**
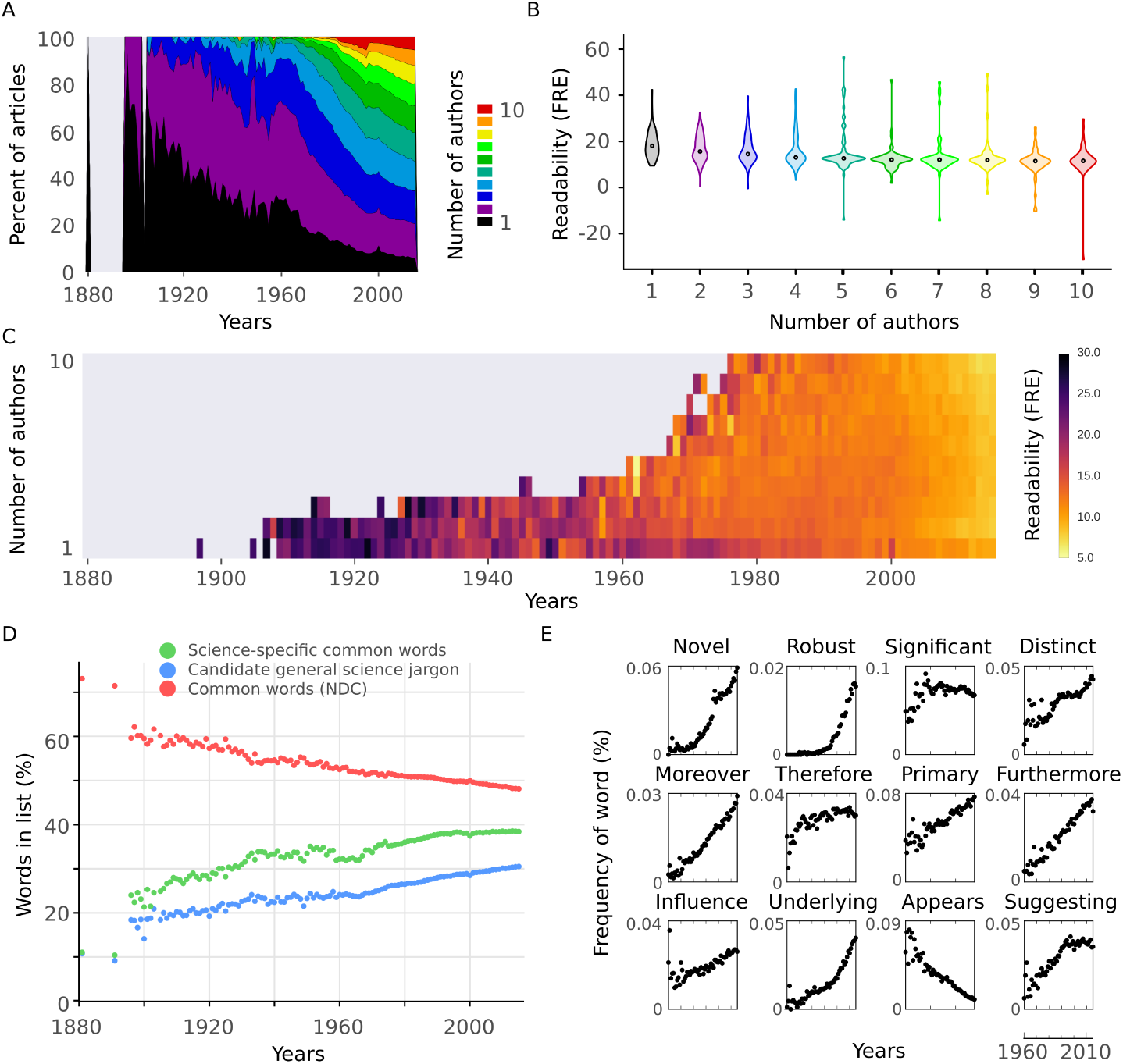
Readability is affected by the number of authors and general scientific jargon. **a**, Proportion of number of authors per year for all articles in the abstract corpus. **b**, Distributions of Flesch Reading Ease (FRE) scores for different numbers of authors (1-10). For New Dale-Chall (NDC), see Supplementary Materials S4, **c**, Mean FRE score for each year for different numbers of authors (1-10). For visualisation purposes, bins with fewer than ten abstracts are excluded. For NDC, see Supplementary Materials S5. **d**, Mean percentage of words in abstracts per year included in three different lists: science-specific common words (green, 2949 words), general scientific jargon (blue, 2140 words) and NDC common words (red, 2949 words). **e**, Example candidate jargon words taken from the general scientific jargon list. Mean percentage of each word’s frequency in abstracts per year is shown.

To test the first hypothesis, we divided the data by the number of authors. More authors were associated with decreased readability (Fig. 3B, Supplementary Materials S4). However, we observed the same trend of decreasing readability across years regardless of the number of authors (Fig. 3C, Supplementary Materials S5). When we included the number of authors as a predictor in the linear mixed effects model, it was found to be significantly related to readability (FRE: *b* = -0.23, *p* < 10^−15^; NDC: *b* = 0.033, *p* < 10^−15^), while the fixed effect of year remained significant (FRE: *b* = -0.17, *p* < 10^−15^; NDC: *b* = 0.014, *p* < 10^−15^, Supplementary Materials S6). We can therefore reject the hypothesis that the increase in the number of authors on scientific articles is responsible for the observed trend, although abstract readability decreases with more authors.

To test the second hypothesis, we constructed a measure for in-group scientific vocabulary. We selected the 2949 most common words which were not included in the NDC common word list from 12 000 abstracts sampled at random (see Methods for procedure). This is analogous to a ‘science-specific common word list’. This list also includes topics which have increased over time (e.g. ‘gene’) and subject-specific words (e.g. ‘tumor’), which are not indicative of an in-group scientific vocabulary. We removed such words to create a general scientific jargon list (2140 words, see Methods and Supplementary Materials S7). While the percentage of common words from the NDC common word list decreased with year (r = -0.93, *p* < 10^−15^, Fig. 3D), there was an increase in the percentage of science-specific common words (r = 0.90, *p* < 10^−15^) and general scientific jargon (r = 0.95, *p* < 10^−15^) (Fig. 3D). Twelve candidate jargon words are presented in Fig. 3E. While one word (‘appears’) decreased with time, all the remaining examples show sharp increases over time. Taken together, this provides evidence in favour of the hypothesis that there is an increase in general scientific jargon which partially accounts for the decreasing readability.

## DISCUSSION

From analysing over 700 000 abstracts in 122 biomedical journals we have shown that a steady decrease of readability over time is present in the scientific literature. It is important to put the magnitude of these results in context. A FRE score of 100 is designed to reflect the reading level of a 10-11 year old. A score between 0 and 30 is considered understandable by college graduates (1, 2). In 1960, 16.3% of the texts in our corpus had a FRE below 0. In 2015, this number had risen to 26.5%. In other words, more than a quarter of scientific abstracts now have a readability considered beyond college graduate level English. We then validated abstract readability against full text readability, demonstrating that it functions as a suitable approximation for comparing main texts.

We investigated two possible reasons why this trend has occurred. First, we found that readability of abstracts correlates with the number of co-authors, but this failed to fully account for the trend through time. Second, we showed that there is an increase in general scientific jargon over years, indicative of a progressively increasing in-group scientific language (“science-ese”).

An alternative explanation for the main finding is that the cumulative growth of scientific knowledge necessitates increasingly complex language. This cannot be directly tested, but if this were to fully explain the trend, we would expect a greater diversity of vocabulary as science grows more specialised. While accounting for the original finding of the increase in difficult words and of syllable count, this would *not* explain the increase of general scientific jargon words (e.g. ‘furthermore’ or ‘novel’, Fig. 3E). Thus, this possible explanation cannot fully account for our findings.

Lower readability implies less accessibility, particularly for non-specialists. Since the 1950’s scientific literacy has been a fundamental goal in the educational sciences, and has even been considered “essential for effective citizenship” (16). Fewer than 30% of American adults are scientifically literate (17, 18). Even more problematic, recent global multi-year measures show stagnating or decreasing trends in scientific literacy in children (19). Our results, combined with the trends in scientific literacy, are worrisome. In addition, amidst concerns that modern societies are becoming less stringent with actual truths, replaced with true-sounding “post-facts” (20, 21) science should be advancing our most accurate knowledge. One way to achieve this is for science to maximise its accessibility to non-specialists such as journalists, policy-makers and the wider public.

Lower readability is also a problem for specialists (22, 23, 24). This was explicitly shown by Hartley (22) who demonstrated that rewriting scientific abstracts, to improve their readability, increased academics’ ability to comprehend them. While science is complex, and some jargon is unavoidable (25), this does not justify the continuing trend that we have shown. It is also worth considering the importance of comprehensibility of scientific texts in light of the recent controversy regarding the reproducibility of science (26, 27, 28, 29, 30). Reproducibility requires that findings can be verified independently. To achieve this, reporting of methods and results must be sufficiently understandable.

What can be done to reverse this trend? Scientists themselves can estimate their own readability in most word processing software. Further, while some journals aim for readability, perhaps a more stringent review of article readability is required during the review process by journals. Finally, in an era of data metrics, it is possible to assess a scientist’s average readability, analogous to the h-index for citations (31). Such an ‘r-index’ could be considered an asset for those scientists who emphasise clarity in their writing.

## MATERIALS AND METHODS

### Journal Selection

We aimed to obtain journals with high impact factor from a representative selection of the life sciences and biomedicine, and which were indexed on PubMed.

Using the Thomson Reuters Research Front Maps (*http://archive.sciencewatch.com/dr/rfm/*) and the Thomson Reuters Journal Citation Reports, we selected 12 fields. From each of eleven of the fields, twelve journals were selected. The final field (Multidisciplinary) only contained six journals. Some journals exist in multiple fields, thus the number of journals (122) is below the possible maximum of 138 journals. See Supplementary Materials S1 for the journals and their field mappings. See Supplementary Methods for the full selection criteria.

Articles were downloaded from PubMed between April 22 2016 and May 15 2016. The text of the abstract, journal name, title of article, PubMed IDs and publication year were extracted. Throughout the article, we only used data up to and including 2015.

### Language preprocessing

Abstracts downloaded from PubMed were preprocessed so that the words and syllables could be counted. TreeTagger (32) was used to identify sentence endings and to remove non-words (e.g. numbers) and any remaining punctuation from the abstracts. Scientific texts contain numerous phrasings which TreeTagger did not parse adequately. We did three rounds of quality control where at least 200 preprocessed articles, sampled at random, were compared with their original texts. After identifying irregularities with the TreeTagger performance, regular expression heuristics were created to prepare the abstracts prior to using the TreeTagger algorithm. After the three rounds of quality control, the stripped abstracts contained only words with at least one syllable and periods to end sentences. Sentences containing only one word were ignored.

The heuristic rules after quality control rounds included: removing all abbreviations, adding spaces after periods when missing, adding a final period at the end of the abstract when missing, removing numbers that ended sentences, identifying sentences that end with “etc.” and keeping the period, removing all single letter words except ‘a’, ‘A’ and ‘I’, removing nucleic acid sequences, replacing hyphens with a space, removing periods arising from the use of binomial nomenclature, and removing copyright and funding information. All preprocessing scripts are available at github.com/wiheto/readabilityinscience. Examples of texts before and after preprocessing are presented in Supplementary Materials S8. We confirmed that the observed trends were not induced by the preprocessing steps by running the readability analysis presented in Fig. 1D,E using the raw data (Supplementary Materials S9).

### Language and Readability metrics

Two well-established readability measures were used throughout the article: the Flesch Reading Ease (FRE) (1, 2) and the New Dale-Chall Readability Formula (NDC) (3). These measures are comprised of different language metrics: syllable count, sentence count, word count and percentage of difficult words.

Counting the syllables of a word was performed in a three step fashion. First, the word was required to have a vowel or a ‘y’ in it. Second, the word was queried against a dictionary that contained specified syllable counts using the natural language toolkit (NLTK (33)). If there were multiple possible syllable counts for a given word, the longer alternative was chosen. Third, if the word was not in the dictionary, the number of vowels (excluding diphthongs) was counted. If a word ended in a ‘y’, this was counted as an additional syllable in this third step.

Word count was calculated by counting all the words in the abstract that had at least one syllable. The number of sentences was calculated by counting the number of periods in the preprocessed abstracts.

The percentage of difficult words originated from (3), defined as words which do not belong to a list of common words. The “NDC common word” list used here was taken from the NDC implementation in the textstat python package (https://github.com/shivam5992/textstat/) which included 2949 words (Supplementary Materials S7). This list excludes some words from the original NDC common word list (such as abbreviations, e.g. ‘A.M.’; and double words, e.g. ‘all right’).

FRE uses both the average number of syllables per word and the average number of words per sentence to estimate the reading level.

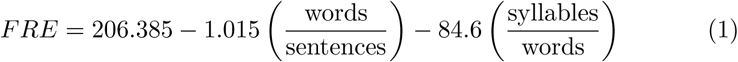
where “words”, “sentences” and “syllables” entail the number of each in the text respectively.

NDC scores are calculated by using the percentage of difficult words and the average sentence length of abstracts. While the NDC was originally calculated on 100 words due to computational limitations, we used the entire text.

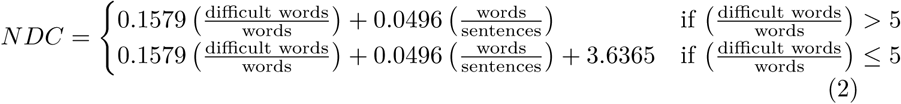
where “words”, and “sentences” entail the number of each in the text respectively. “difficult words” is the number of words that are not present in the NDC common word list.

We have used two well-established readability formulae in our analysis. It is important to consider readability formulae as an estimate of readability (5, 4) and not a direct measure of absolute understandability of a text. The application of readability formulae has been questioned (34, 35, 36) and modern alternatives have been proposed (see 37). However, NDC has been shown to perform comparably with these more modern methods (37). Text from different media can be underestimated or overestimated in their ‘absolute’ readability levels based on other factors, such as the overall coherence of a text (4, 36).

However, this is more of a problem when comparing readability between text media, than when examining trends of readability over time within a specific medium, such as scientific texts.

### Science-specific common word and candidate general science jargon lists

To construct the science-specific common word list, 12 000 articles were selected to identify words frequently used in the scientific literature. In order to avoid any recency bias, 2000 articles were randomly selected from six different decades (starting at the 1960s). From these articles, the frequency of all words was calculated. After excluding words in the NDC common word list, the 2949 most frequent words were selected. The number 2949 was selected to be the same length as the NDC common word list. This list is the “science-specific common word” list.

To validate that this list is capturing general scientific terminology, we created a verification list by performing the same steps as above on an additional set of 12 000 articles. Of the 2949 words in the science-specific common word list, 91.52% of the words were present in the verification list (see Supplementary Materials S7 for both word lists).

A subset of the science-specific common words was selected to form the “general scientific jargon” list (2140 words in total). This subset was created by filtering the science-specific common word list to leave only candidate jargon words. These words were deemed to reflect those which scientists may use regardless of their field or topic of study. This candidate jargon was identified manually (see Supplementary Methods for identification process and S7 for the list of candidate jargon).

The 24 000 articles used in the derivation and verification of the lists were excluded from all further analysis where they were used.

### Comparison of full texts vs abstracts

To compare the readability of full texts and abstracts, we chose six representative journals from the PubMed Central Open Access Subset for which all full texts of articles were available under a Creative Commons or similar license. These journals were BMC Biology, eLife, Genome Biology, PloS Biology, PloS Medicine and PloS ONE. None of these journals were a part of the original journal list which was used in the main analysis.

In total, 143 958 articles were included in the analysis. Both article abstracts and full texts were preprocessed according to the procedure outlined above and the readability measures were calculated.

## Statistics

All statistical modelling was performed in R version 3.3.0

We evaluated the relationship between the readability of single abstracts and year of publication separately for FRE and NDC scores. The data can be viewed as hierarchically structured since abstracts belonging to different journals may differ in key aspects. In addition, journals span over different ranges of years (Fig. 1C and Supplementary Materials S1). In order to account for this structure, we performed linear mixed effect modelling using the R-packages lme4 version 1.1-12 (38), and lmerTest version 2.0-33 (39) with maximum likelihood estimation. We compared different models of increasing complexity. The included models were as following: (M0) a null model in which readability score was predicted only by journal as random effect with varying intercepts; (M1) the same as M0, but with an added fixed effect of time; and (M2) the same as M1, but with varying slopes for the random effect of journal (Supplementary Materials S2). We selected the best fitting model as determined by the Akaike Information Criterion and the Bayesian Information Criterion.

In order to test that the trend was not explained by the increasing number of authors with year, we specified an additional model. It was identical to M2 above, but also included the number of authors as a second fixed effect. Some articles (n = 2325) lacked author information, and were excluded from the analysis. This model was performed using two sets of the data: i) a subset including only articles with one to ten authors (n = 650 344), ii) a full dataset consisting of all articles with complete author information (n = 705 127) (see Supplementary Materials S6). The motivation for (i) was that abstracts with many authors may bias the results.

## Acknowledgements

This article has a FRE score of 44.5.

## Supplementary Methods

### Details regarding journal selection

#### Fields

First, we identified general fields of scientific study from the life sciences and biomedicine using the Thomson Reuters Research Front Maps (RFM). The Thomson Reuters Journal Citation Reports (JCR) also contain fields, but these fields are more specific and more numerous. We thus identified subfields from the Thomson Reuters Journal Citation Reports (JCR). In order to account for discrepancies between these lists, the RFM fields were altered from having categories of “Biology and Biochemistry” and “Molecular Biology and Genetics” to “Biology” and “Molecular Biology, Genetics and Biochemistry”. This was because the JCR subfield categories contained “Biochemistry and Molecular Biology” as a single subfield. We also added two additional fields: “Multidisciplinary” and “All”. “Multidisciplinary” accounts for journals which publish work from multiple fields, but which did not fit into any one category. “All” accounts for the highest impact journals across all fields.

The final fields consisted of the following:

1. All
2. Biology & Biochemistry
3. Clinical Medicine
4. Immunology
5. Microbiology
6. Molecular Biology, Genetics & Biochemistry
7. Multidisciplinary
8. Neuroscience & Behavior
9. Pharmacology & Toxicology
10. Plant & Animal Science
11. Psychiatry & Psychology
12. Social Sciences, general

From each RFM field, we selected between one and six JCR subfields, deemed to fall within the content of each field.

#### Selection of Journal within Fields

From each field, we selected 12 journals. Journals were semi-automatically selected by querying the PubMed API using R and the package RISmed (1) according to the following criteria:

1. There should be more than 15 years between the years of the first PubMed entries and the most recent PubMed entries. This was done by querying the dates of the minimum year of the first five entries, and the maximum year of the most recent five entries.
2. There should be more than 100 articles listed on PubMed for the journal.
3. The impact factor of the journal should not be below 1 according to the 2015 Thomson Reuters Journal Citation Report.
4. The articles within the journal should be in English.

Due to a lack of journals which fulfilled the fourth criterion for the Multidisciplinary field, we restricted this field to only 6 journals.

The selection of journals from fields was conducted according to the following criteria in order of importance.

1. The journals from each RFM field should be selected as equally as possible from each of the constituent JCR subfields.
2. The journals should be selected by highest impact factor within each subfield after fulfilling the above criterion.
3. When the same journal exists in more than one JCR subfield of the same RFM field, the next journal should be taken from that JCR subfield such that list sizes are as equal as possible.
4. When conflicts arise regarding which JCR subfield a journal should be selected from, the journal with the higher impact factor should be selected.

The final journal list is presented in Supplementary Materials S1.

1. Stephanie Kovalchik (2016). RISmed: Download Content from NCBI Databases. R package version 2.1.6. https://CRAN.R-project.org/package=RISmed

### Identifying candidate jargon by manually filtering the science common word list

The science-specific common word list consisted of 2949 words (see Methods section). This was filtered down to 2140 words which contained candidate jargon.

All four co-authors went through the list by themselves. Prior to going through the list the following guidelines were formulated:

1. Is the word unwanted for some other reason (e.g. abbreviations or units that survived the preprocessing (e.g. “mmol”))˙.
2. Does the word refer to or used within a field specific subfield (e.g., “hepatitis”).
3. Is the word a general word or area of study which could be increasing in time due to it becoming studied more over time (e.g. “gene”).
4. Is the word a noun, adjective or verb which can be assigned to non-science objects and could instead be part of an easy word list (e.g “mouse”, “green”, “September”).

If not considered any of the above, the word was marked as a possible jargon word (i.e. candidate jargon).

The intention was that the ratings would be performed independently. However, all authors considered this a hard task for two major reasons: (1) many words could be subject specific words or candidate jargon depending on context (e.g. “replication” and “binding” will often be used in subject specific contexts); (2) different authors seemed to have different intuitions about what constituted a word that should be part of a general easy word list. For this reason, there was a consultation between all authors mid-review to control that the guidelines were being performed in a similar way. In this consultation examples of what each author had rated were discussed. Due to this meeting, the ratings can not be classed as completely independent.

It was decided that if 3 out of the 4 authors classified the word as candidate jargon, it would get included on the jargon list (2083 words). Furthermore, the words where 2 authors agree considered the words to be jargon (258 words) were reviewed again collectively. From this list 57 of these words were agreed upon to be included in the jargon list. This meant that 2140 words were in the jargon list. The list of jargon words can be found in Supplementary Materials S08.

### Supplementary materials for “The readability of scientific texts is decreasing over time”

#### Contents

S01 Journal information (tabular *S01_JournalSelection.xlsx*, external document not included in pdf*).

S02 Linear mixed effect model (table).

S03 New Dale-Chall abstracts and full text (figure).

S04 New Dale-Chall for different number of authors (figure).

S05 New Dale-Chall for different number of authors over years (figure).

S06 Linear mixed effect model including number of authors (table).

S07 Word lists (NDC and science-specific common word) (tabular *S09_Wordlists.xlsx*, external document not included in pdf*).

S08 Examples of before/after preprocessing of scientific language (text).

S09 Mean readability over years with only minimal preprocessing (figure).

*see github.com/wiheto/readabilityinscience

**Supplementary Materials S2:** Linear mixed effect model. Model fits for two different linear mixed effect models examining the relationship between readability scores and year. A null model without year as a predictor is included as a baseline comparison. Lower dAIC and dBIC values indicate better model fit.

| Metric | Model | dAIC | dBIC | b | t | df | p |
| --- | --- | --- | --- | --- | --- | --- | --- |
| FRE | M0 | 15542 | 15507 | - | - | - | - |
| | M1 | 5134 | 5111 | -0.144 | -102.4 | 707413 | $p < 10^{-15}$ |
| | M2 | 0 | 0 | -0.189 | -12.5 | 122 | $p < 10^{-15}$ |
| NDC | M0 | 28550 | 28516 | - | - | - | - |
| | M1 | 4078 | 4055 | 0.014 | 157.8 | 707435 | $p < 10^{-15}$ |
| | M2 | 0 | 0 | 0.016 | 20.4 | 116 | $p < 10^{-15}$ |

FRE = Flesch Reading Ease; NDC = New Dale-Chall Readability Formula; M0 = Journal as random effect with varying intercepts; M1 = M0 with an added fixed effect of time; M2 = M1 with varying slopes for the random effect of journal; dAIC = difference in Akaike Information Criterion from the best fitting model (M2); dBIC = difference in Bayesian Information Criterion from the best fitting model (M2); df = Degrees of Freedom calculated using Satterthwaite approximation.

**Supplementary Materials S3:**
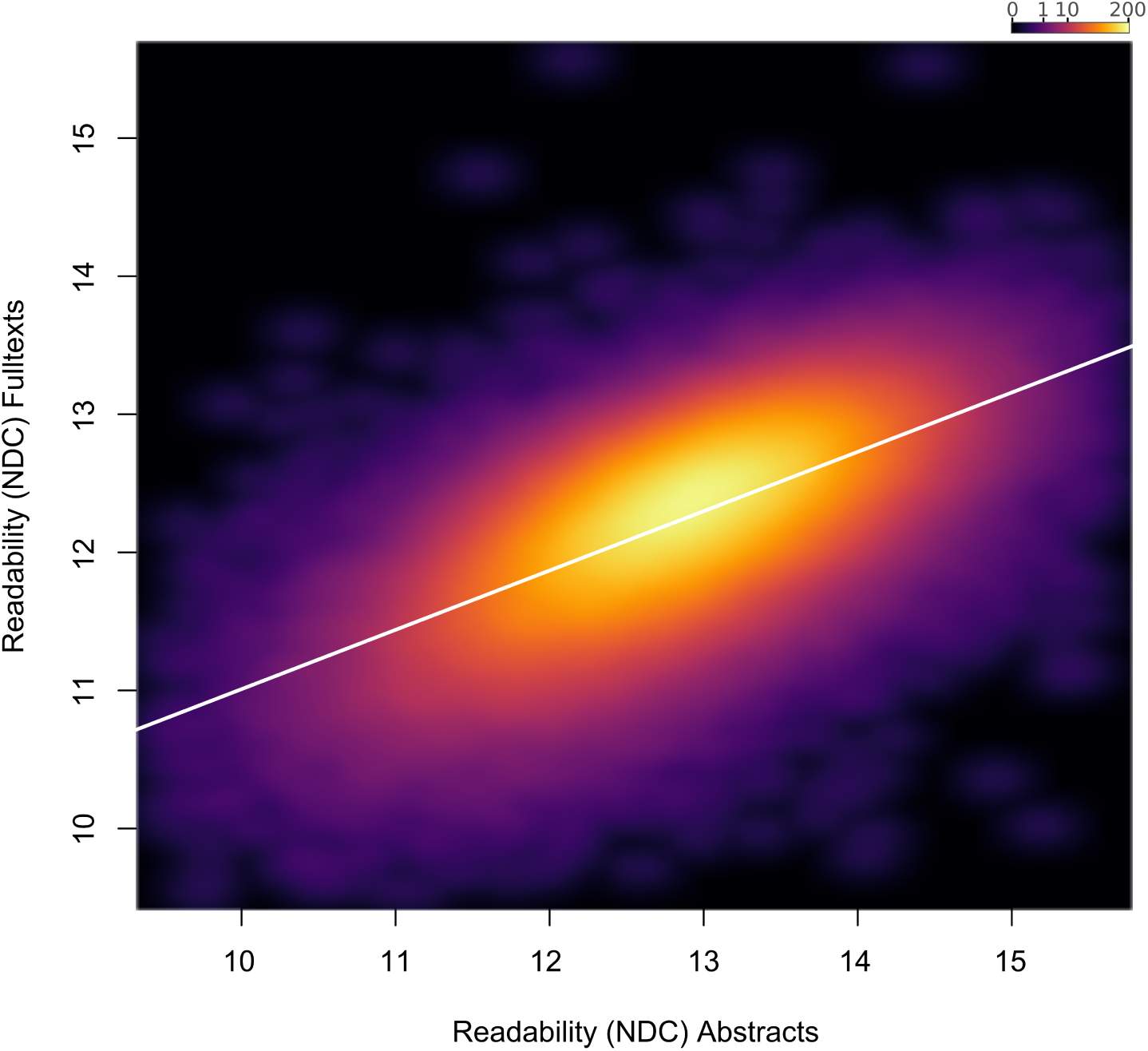
New Dale-Chall abstracts and full text. Relationship between New Dale-Chall Readability Formula scores of abstracts and full texts across the full text corpus, depicted by a two-dimensional kernel density estimate. Axis limits are set to include at least 99% of the data.

**Supplementary Materials S4:**
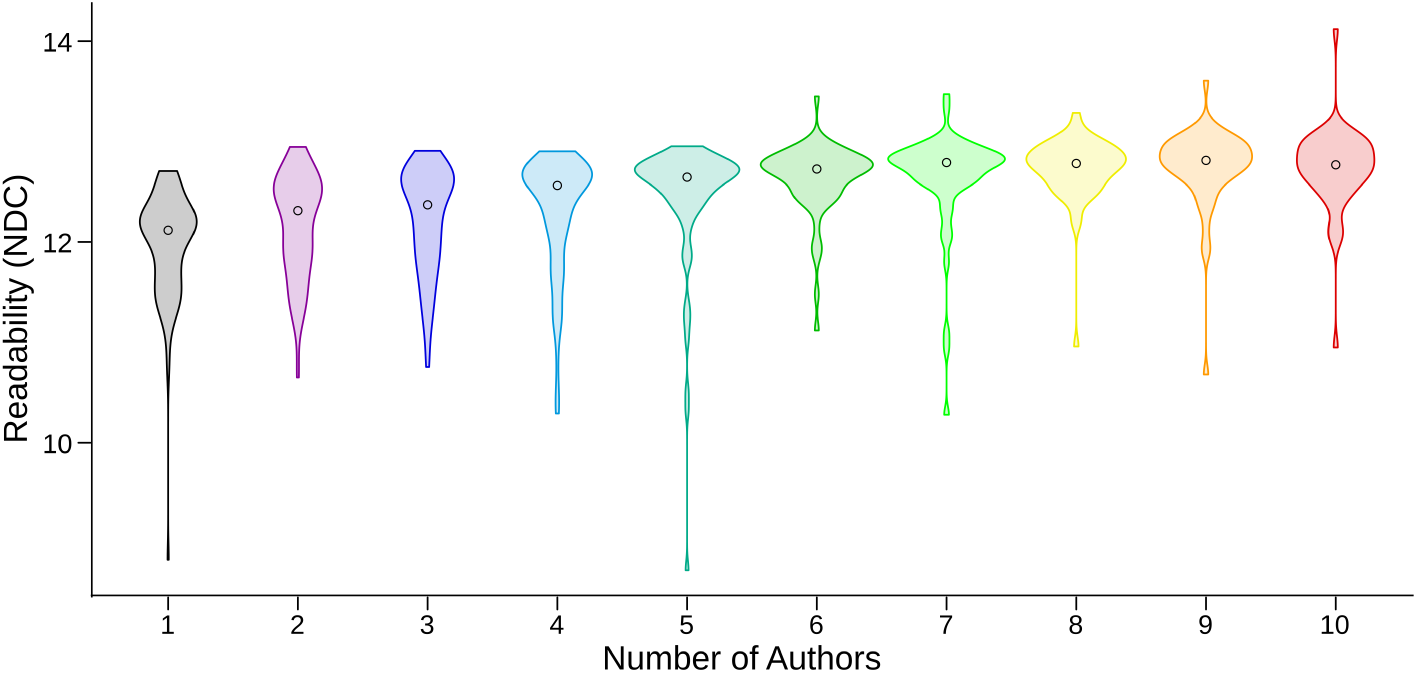
New Dale-Chall for different number of authors. Distributions of New Dale-Chall Readability Formula scores for different numbers of authors (1-10). Higher scores indicate less readability.

**Supplementary Materials S5:**
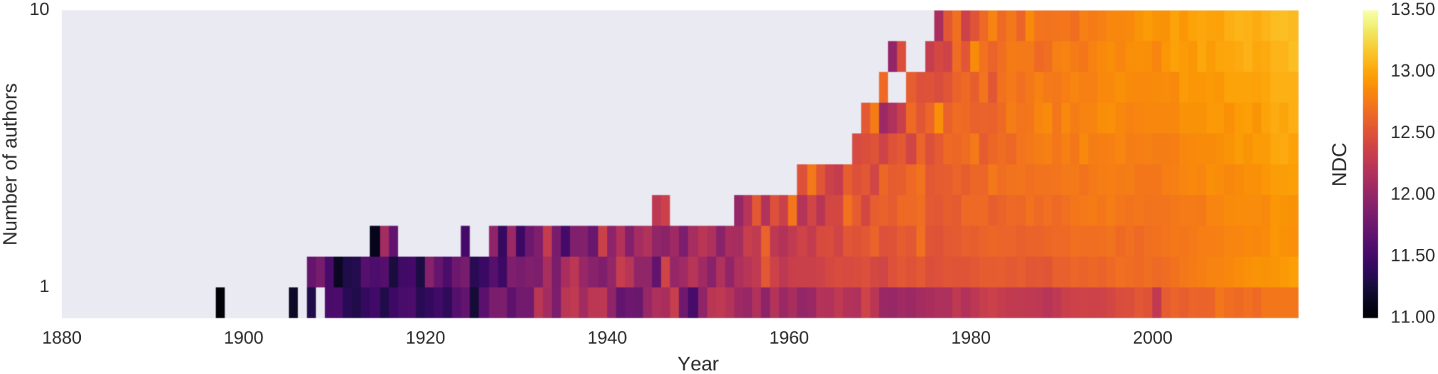
New Dale-Chall for different number of authors over years. Mean New Dale-Chall Readability Formula score for each year for different numbers of authors (1-10). For visualisation purposes, bins with fewer than ten abstracts are excluded. Higher scores indicate less readability.

**Supplementary Materials S6:**
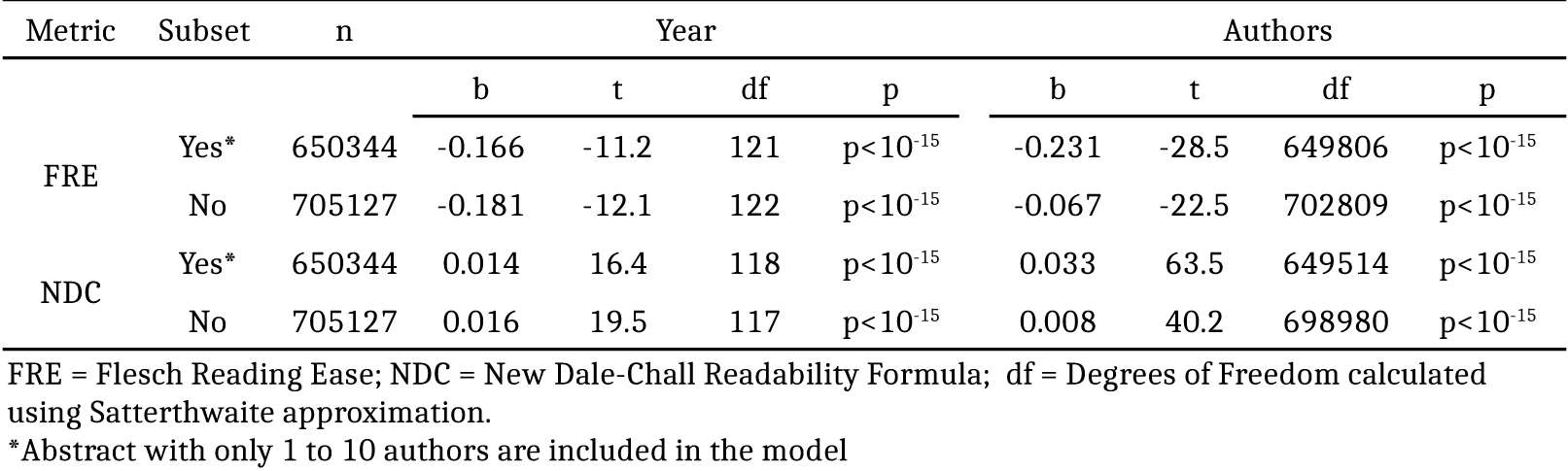
Linear mixed effect model including number of authors. Linear mixed effect models predicting readability scores by year and number of authors with journals as random effect.

| Metric | Subset | n | Year |  |  |  | Authors |  |  |  |
| --- | --- | --- | --- | --- | --- | --- | --- | --- | --- | --- |
|  |  |  | b | t | df | p | b | t | df | p |
| FRE | Yes* | 650344 | -0.166 | -11.2 | 121 | $p < 10^{-15}$ | -0.231 | -28.5 | 649806 | $p < 10^{-15}$ |
| | No | 705127 | -0.181 | -12.1 | 122 | $p < 10^{-15}$ | -0.067 | -22.5 | 702809 | $p < 10^{-15}$ |
| NDC | Yes* | 650344 | 0.014 | 16.4 | 118 | $p < 10^{-15}$ | 0.033 | 63.5 | 649514 | $p < 10^{-15}$ |
| | No | 705127 | 0.016 | 19.5 | 117 | $p < 10^{-15}$ | 0.008 | 40.2 | 698980 | $p < 10^{-15}$ |

FRE = Flesch Reading Ease; NDC = New Dale-Chall Readability Formula; df = Degrees of Freedom calculated using Satterthwaite approximation.
\*Abstract with only 1 to 10 authors are included in the model

**Supplementary Materials S8: Examples of before/after preprocessing of scientific language.**

In this document there are 8 examples where we quote the original abstract text, followed by the output from the preprocessing steps which removes many unwanted features of text for quantifying readability. The articles have been selected to demonstrate the effectiveness of the preprocessing steps across disciplines and year of publication.

These examples are omitted in the preprint edition due to possible copywrite concerns of quoting entire abstracts from journals

**Supplementary Materials S9:**
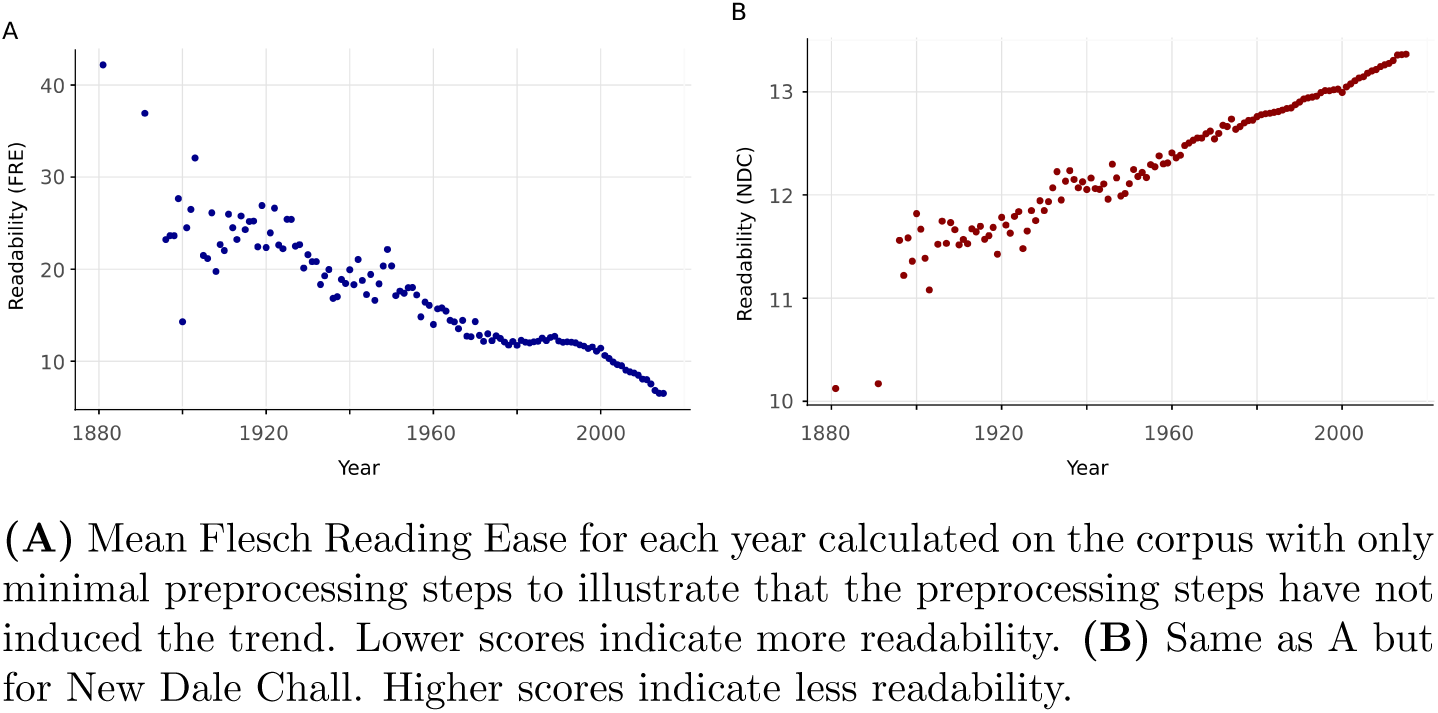
Readability over years with minimal preprocessing. (**A**) Mean Flesch Reading Ease for each year calculated on the corpus with only minimal preprocessing steps to illustrate that the preprocessing steps have not induced the trend. Lower scores indicate more readability. (**B**) Same as A but for New Dale Chall. Higher scores indicate less readability.

